# Colorectal cancer metastases in the liver establish immunosuppressive spatial networking between tumor associated *SPP1*^+^ macrophages and fibroblasts

**DOI:** 10.1101/2020.09.01.273672

**Authors:** Anuja Sathe, Kaishu Mason, Susan M. Grimes, Zilu Zhou, Billy T. Lau, Xiangqi Bai, Andrew Su, Xiao Tan, HoJoon Lee, Carlos J. Suarez, Quan Nguyen, George Poultsides, Nancy R. Zhang, Hanlee P. Ji

## Abstract

**Purpose:** The liver is the most frequent metastatic site for colorectal cancer (**CRC**). Its microenvironment is modified to provide a niche that allows CRC cell growth. This study focused on characterizing the cellular changes in the metastatic CRC (**mCRC**) liver tumor microenvironment (**TME**).

**Experimental Design:** We analyzed a series of microsatellite stable (MSS) mCRCs to the liver, paired normal liver tissue and peripheral blood mononuclear cells using single cell RNA-seq (**scRNA-seq**). We validated our findings using multiplexed spatial imaging and bulk gene expression with cell deconvolution.

**Results:** We identified TME-specific *SPP1*-expressing macrophages with altered metabolism features, foam cell characteristics and increased activity for extracellular matrix (**ECM**) organization. *SPP1+* macrophages and fibroblasts expressed complementary ligand receptor pairs with the potential to mutually influence their gene expression programs. TME lacked dysfunctional CD8 T cells and contained regulatory T cells, indicative of immunosuppression. Spatial imaging validated these cell states in the TME. Moreover, TME macrophages and fibroblasts had close spatial proximity, a requirement for intercellular communication and networking. In an independent cohort of mCRCs in the liver, we confirmed the presence of *SPP1*^+^ macrophages and fibroblasts using gene expression data. An increased proportion of TME fibroblasts was associated with worst prognosis in these patients.

**Conclusions:** We demonstrated that mCRC in the liver is characterized by transcriptional alterations of macrophages in the TME. Intercellular networking between macrophages and fibroblasts supports CRC growth in the immunosuppressed metastatic niche in the liver. These features can be used to target these immune checkpoint resistant MSS tumors.

**TRANSLATIONAL RELEVANCE:** The liver is the commonest site for metastatic colorectal cancer (**mCRC**). Alterations in the tumor microenvironment (**TME**) allow metastatic cells to seed the distant liver site and grow. Leveraging single-cell RNA sequencing, we discovered a distinct *SPP1*+ macrophage cell state with pro-fibrogenic gene expression and altered metabolism. These *SPP1*+ macrophages communicated with fibroblasts, mutually influencing each other’s gene expression program. Using spatial imaging, we confirmed proximal colocalization between macrophages and fibroblasts in the mCRC TME, which is required for intercellular communication. These states and intercellular communication promoted immunosuppression in the TME, with a lack of dysfunctional anti-tumor CD8 T cells and prevalence of regulatory T cells. Increased fibroblasts were associated with worst prognosis in an independent patient cohort. Our results identified novel TME features that result in reshaping of the metastatic niche that allows progression of mCRC. These features can be potential targets for mCRC treatment, which is microsatellite stable and resistant to immune checkpoint blockade.

## INTRODUCTION

Nearly 50% of all patients with colorectal cancer (**CRC**) have metastases. The most common site for metastatic colorectal cancer (**mCRC**) is the liver (1). Through hematogenous spread, colon cancer cells reach the liver and establish themselves in the hepatic parenchyma. Liver metastasis is a major contributor to morbidity and mortality of Stage IV patients. There are specific cellular processes that enable a CRC metastasis to establish itself in the liver. The hepatic microenvironment and its cellular composition are functionally quite different than those present within the colon microenvironment encompassing the primary tumor. The various cells in the liver microenvironment must be altered to accommodate foreign colon cancer cells (2). Cellular changes specific to the liver facilitate mCRCs growth and play a role in suppressing the patient’s immune response (3, 4).

Immune checkpoint blockade targets the TME-based T-cells and their communication with tumors cells via PD-1/PD-L1. Only mCRCs with microsatellite instability (**MSI**), a hypermutable state, respond to immunotherapy. However, only around 4% of mCRC tumors are of the MSI subtype (5). Majority of mCRCs are microsatellite stable (**MSS**) and show no response to checkpoint blockade. Furthermore, MSS mCRC tumors generally do not have significant levels of T cell infiltration which is a requirement for effective checkpoint blockade (3, 6). As a result, MSS mCRCs have a profoundly immunosuppressed TME and are highly resistant to immunotherapy. To develop effective immune-based therapeutic strategies for mCRC, it is essential to characterize the cell states and interactive networking present within the TME in sites such as the liver.

Metastatic colorectal cancers in the liver are a complex mixture of many different cell types originating from the tumor epithelium, immune system, and hepatic stroma. Even for a specific cell type, there are different cell “states” reflecting functional variation depending on the local cellular context as seen in liver parenchyma. Several recent studies have used single-cell RNA (**scRNA-seq**) to characterize the various cell types and the functional states in the CRC TME in colon. These studies represent a survey of the primary colon tumor. There are only a few single cell genomic studies of mCRCs in the liver. These scRNA-seq studies have focused on the immune cells isolated from the mCRC TME (4, 7, 8). However, non-immune cell types such as fibroblasts and endothelial cells also contribute to the mCRC TME. Intercellular signaling and networking among the immune and non-immune cells orchestrate metastatic progression (9).

We conducted a study to determine the multi-cellular features and interactions for mCRCs in the liver. Our analysis specifically focused on MSS mCRCs. We used a multi-pronged approach: (1) single-cell RNA (**scRNA-seq**); (2) spatial multiplexed imaging; (3) conventional RNA-seq. For the single cell studies, the tumors were analyzed directly without flow sorting – this approach preserved the composition of the native cells in the liver metastasis. We identified a unique category of TME-based macrophages that networks with cancer associated fibroblasts (**CAF**). We examined the spatial organization and proximity of these and other cell types with multiplexed imaging. Our results showed close proximity of macrophages and CAFs compared to other TME cell types. Using an independent set of mCRCs in the liver with RNA-seq data, cellular deconvolution identified these different cell types and confirmed our single cell results.

## MATERIALS AND METHODS

### Sample collection and processing

This study was conducted in compliance with the Helsinki Declaration. All patients were enrolled according to a study protocol approved by the Stanford University School of Medicine Institutional Review Board (IRB-11886 and IRB-44036). Informed consent was obtained from all patients.

Tissue samples came from surgical resections or matched normal tissue from sites displaced at least several centimeters from the tumor. Tissues were collected in plain RPMI on ice immediately after resection and dissected with iris scissors. Single-cell suspensions were obtained from tissue fragments using enzymatic and mechanical dissociation and from peripheral blood using peripheral blood mononuclear cell (**PBMC**) isolation as described previously (10). Briefly, cells were washed twice in RPMI + 10% FBS, filtered through 70 μm (Flowmi, Bel-Art SP Scienceware, Wayne, NJ), followed by 40 μm filter (Flowmi). Cryofrozen cells were rapidly thawed in a bead bath at 37 °C followed by above washing and filtering steps. Live cell counts were obtained on a BioRad TC20 cell counter (Biorad, Hercules, CA) or a Countess II FL Automated Cell Counter (ThermoFisher Scientific) using 1:1 trypan blue dilution. Cells were concentrated between 500-1500 live cells/μl for scRNA-seq.

### Histopathology

Tissue was fixed in 10% formalin for approximately 24 hours at room temperature. Paraffin embedding and hematoxylin and eosin staining was conducted by the Human Pathology Histology Services core facility at Stanford University. We reviewed clinical histopathology reports for all patients that examined the expression of HER2, special stains for MLH1, MSH2, MSH6 and PMS2 for MSI/MSS status using standard clinical immunohistochemistry (**IHC**) protocols.

### Single-cell RNA sequencing

The scRNA-seq libraries were generated from cell suspensions using Chromium Single Cell 3′ Library & Gel Bead Kit v2 or Chromium Next GEM Single cell Immune Profiling 5’ v1.1 (for P8640) (10X Genomics, Pleasanton, CA, USA) as per manufacturer’s protocol and sequenced on Illumina sequencers (Illumina, San Diego, CA). All libraries from a patient were prepared in the same experimental batch. Ten thousand cells were targeted from tissue dissociation suspensions and 3000 for PBMCs with 14 PCR cycles for cDNA and library amplification. A 1% or 2% E-Gel (ThermoFisher Scientific, Waltham, MA, USA) was used for quality control evaluation of intermediate products and sequencing libraries. A Qubit (Thermofisher Scientific) or qPCR with Kapa library quantification kit (Kapa Biosystems, Wilmington, MA) was used to quantify the libraries as per the manufacturer’s protocol.

### Processing scRNA-seq data

Cell Ranger (10x Genomics) version 3.1.0 ‘mkfastq’ and ‘count’ commands were used with default parameters and alignment to GRCh38 to generate matrix of unique molecular identifier **(UMI)** counts per gene and associated cell barcode. We constructed Seurat objects from each dataset using Seurat (version 4.0.1) (11, 12) to apply quality control filters. We removed cells that expressed fewer than 200 genes, had greater than 30% mitochondrial genes or had UMI counts greater than 8000 which is an indicator of cell doublets. We removed genes that were detected in less than 3 cells. We normalized data using ‘SCTransform’ and used first 20 principal components with a resolution of 0.6 for clustering. We then removed computationally identified doublets from each dataset using DoubletFinder (version 2.0.2) (13). The ‘pN’ value was set to default value of 0.25 as the proportion of artificial doublets. The ‘nExP’ was set to expected doublet rate according to Chromium Single Cell 3’ v2 reagents kit user guide (10x Genomics). These parameters were used as input to the ‘doubletFinder_v3’ function with number of principal components set to 20 to identify doublet cells.

### Batch-corrected integrated scRNA-seq analysis

Individual Seurat objects were merged and normalized using ‘SCTransform’ (11, 12). To eliminate potential batch effects, we integrated all datasets across experimental batches by using a soft variant of k-means clustering implemented in the Harmony algorithm (version 0.1.0) (14). The experimental batch metrics were used in the grouping variable in the ‘RunHarmony’ function, and this reduction was used in both ‘RunUMAP’ and ‘FindNeighbors’ functions for clustering. The first 20 principal components and a resolution of 1 was used for clustering. We used the Adjusted Rand Index (**ARI**) to compare similarity between cluster labels and experimental batch meta data label for each cell. A vector of these respective class labels was supplied to the ‘adjustedRandIndex’ function in mclust package (v 5.4.7) (15). The data from the ‘RNA’ assay was used for all further downstream analysis with other packages, gene level visualization or differential expression analysis. The data was normalized to the logarithmic scale and the effects of variation in sequencing depth were regressed out by including ‘nCount_RNA’ as a parameter in the ‘ScaleData’ function. Differential gene expression analysis was conducted using the ‘FindAllMarkers’ or ‘FindMarkers’ functions respectively using Wilcoxon rank sum test. Parameters provided for these functions were as follows: genes detected in at least 25% cells and differential expression threshold of 0.25 log fold change. Significant genes were determined with *p* < 0.05 following Bonferroni correction. The ‘DoHeatmap’, ‘FeaturePlot’, ‘DimPlot’, ‘DotPlot’, ‘VlnPlot’ functions were used for visualization.

### Cell lineage identification and reclustering of integrated scRNA-seq data

From the batch-corrected Seurat object, cell lineages were identified based on marker gene expression. Red blood cell and platelet clusters were filtered out from the downstream analysis. A single proliferative cluster containing both epithelial and T cells was split based on the expression of normalized counts for *EPCAM* > 0 in epithelial cells. We performed a secondary clustering analysis of each lineage with integration across experimental batches using Harmony and a cluster resolution of 0.6. Any clusters identified as belonging to another cell lineage were united with their lineage counterparts for a second clustering run. This yielded final lineage-specific re-clustering results. In lymphocyte re-clustering, a single cluster containing naïve CD4 and CD8 T cells was gated for CD8 T cells based on the expression of normalized counts for *CD8A* or *CD8B* >0.

### Pathway analysis

Differentially expressed genes in tumor macrophages were used as input to pathway analysis using ‘Reactome_2016’ in the package enrichR (v2.1) (16). We used the ‘AddModuleScore’ function in Seurat to calculate the average expression of a custom gene set of interest. Using this function, genes of interest were first binned into 24 bins of expression levels based on their average expression. From each bin, control genes were randomly selected using default parameters used in this function. Finally, average expression score was calculated as the difference between average expression of gene set of interest and average expression of control genes. Expression between clusters was compared using t-test. Gene signatures of scar associated macrophages from liver cirrhosis (17) and atherosclerotic foam cells (18) were obtained from the original publications. CD8 cytotoxicity signature (*GZMA*, *GZMB*, *GZMK*, *GZMH*, *GNLY*, *PRF1*, *IFNG*, *NKG7*, *KLRK1*, *KLRB1*, *KLRD1*, *CTSW*, *CST7*, *CCL4*, *CCL3*) was compiled from previous publications (19, 20).

### Copy number analysis

InferCNV (version 1.2.3) (21) was used to infer large-scale copy number variations in tumor epithelial cells. As a reference control, we used all myeloid and stromal cells from tumor and normal samples. Count data was used as input. Filtering, normalization and centering by normal gene expression were performed using default parameters. A cut-off of 0.1 was used for the minimum average read counts per gene among reference cells. An additional denoising filter was used with a threshold of 0.2. Copy number variation was predicted using the default six state Hidden Markov Model.

### Receptor-ligand communication between cell types

We obtained the expression matrix from tumor samples using the ‘data’ slot of the ‘RNA’ assay following lineage-specific secondary clustering analysis. We excluded epithelial cells from P6198 with neuroendocrine differentiation from this analysis. This expression matrix was used as input to CellChat (v0.5.0) (22). ‘CellChatDB.human’ was used as the receptor-ligand interaction database. ‘identifyOverExpressedGenes’ and ‘identifyOverExpressedInteractions’ functions were used to identify over-expressed ligands, receptors and interactions in each cell group. Number of interactions were calculated using the ‘aggregateNet’ function and visualized using ‘netVisual_circle’.

We also predicted receptor-ligand interactions likely to affect specific gene expression changes in a target cell lineage using nichenetr (v0.1.0) (23). This analysis utilizes ligand-target regulatory potential scores calculated from prior information. We performed this analysis on fibroblasts as target cells for genes that overlapped with the matrisome program (identified in Supplemental Table 6). We also examined macrophages as target cells with genes belonging to enriched Reactome pathways: ‘Metabolism’, ‘Degradation of the extracellular matrix’, ‘Extracellular matrix organization’, ‘Collagen degradation’ (identified in Supplemental Table 5). NicheNet’s prior models and networks were obtained from https://zenodo.org/record/3260758#.X0WX7BNKhTY.

Ligands predicted to influence expression of genes of interest in target population were calculated using the function ‘predict_ligand_activities’ with default parameters that outputs activity as Pearson correlation coefficient based on prior modelling. The weight or inferred regulatory score between a target gene and ligand was obtained using ‘get_weighted_ligand_target_links’ function. Top 20 ligands and interactions with regulatory potential value in the top 60% were used for visualization. Ligands were assigned to a particular cell type as sender if their expression was greater than one standard deviation from the average ligand expression. Target genes and ligands were visualized using the ‘chordDiagram’ function from the circlize R package (v0.4.11) with transparency scaled to respective regulatory potential value.

### EcoTyper analysis for cell state discovery

Cell state discovery on scRNA-seq expression data was performed using EcoTyper (24) using the scripts and vignette provided on https://github.com/digitalcytometry/ecotyper. The number of NMF restarts was set to 50 and maximum number of states per cell type was set to 10.

### RNA-seq analysis and cell type deconvolution

Fastq files from RNA sequencing of 93 mCRC FFPE samples were obtained from European Genome-Phenome Archive, dataset ID EGAD00001004111 (3). Information on prognosis sub-group for each patient was obtained from the contributors of the data. Data was aligned to genome reference GRCh38 using STAR (v2.6.0a) and transcripts per gene were counted using htseq-count method from HTSeq (v0.5.4). Counts were converted to Transcripts Per kilobase Million (**TPM**) to normalize for gene length, and non-protein-coding genes were removed. Gene length used for normalization was the number of bases covered at least once for all exons in that gene. The TPM value was obtained by calculating the reads per kilobase (**RPK**) for each gene, then calculating the scaling factor as sum (RPK)/10E6 and lastly calculating TPM per gene as RPK/scaling factor. In cases with duplicate sequencing runs for the same patient, TPM counts were averaged.

From our mCRC samples, we obtained the single-cell expression matrix for each cell type using the ‘counts’ slot of the ‘RNA’ assay of the Seurat object with filtering as outlined above. These cell-type gene lists were used as input to CIBERSORTx (25). The signature matrix was created in custom analysis mode using default parameters with minimum expression set to zero and was used for cell fraction imputation. TPM counts from bulk expression dataset was used as the mixture file. Default parameters were used except quantile normalization was disabled, permutations for significance analysis were set to 1000 and batch correction was applied in ‘S-mode’. Resulting proportions were recalculated as a fraction of only TME lineages by removing epithelial cells. Patients were grouped according to their sub-group for overall survival and significant differences in proportion of cell types were assessed by ANOVA with Tukey HSD correction.

### CODEX staining and imaging

A custom antibody panel was developed and validated (Enable Medicine, Menlo Park, CA, USA) for multiplexed imaging with co-detection by indexing (**CODEX**) (Akoya Biosciences, Menlo Park, CA, USA). This imaging technique utilizes antibodies conjugated to unique DNA oligonucleotide barcodes. The CODEX antibodies were validated on formalin fixed paraffin embedded (**FFPE**) tonsil sections and staining patterns were confirmed via comparison with online databases (The Human Protein Atlas, www.proteinatlas.org; Pathology Outlines, www.pathologyoutlines.com) and the published literature. Between 4-6 formalin fixed samples were paraffin embedded into the same tissue block, sectioned at 7 μm, and placed on 22×22 mm glass coverslips (Electron Microscopy Sciences, # 72204-01) pre-coated with poly-L-lysine (Sigma, # P8920). The FFPE tissues on coverslips were stored in a 6-well plate containing storage buffer at 4°C until CODEX acquisition.

CODEX imaging was done as per the manufacturer’s protocol (Akoya Biosciences). Briefly, FFPE tissue sections on coverslips were pretreated by heating on a slide warmer for 25 minutes at 55 degrees C. Tissue deparaffinization and hydration were next performed by incubating the FFPE tissue sections on coverslips for 5 minutes each following a solvent series (Histochoice Clearing Agent, Histochoice Clearing Agent, 100% Ethanol, 100% Ethanol, 90% Ethanol, 70% Ethanol, 50% Ethanol, 30% Ethanol, ddH20, ddH20). Antigen retrieval was performed in 0.01M Citrate Buffer at high pressure. The tissue was washed and equilibrated before staining for 3 hours at room temperature with the 28-plex CODEX antibody cocktail in a staining buffer containing blocking solution (Akoya Biosciences). After staining, the tissues were washed and fixed in 1.6% PFA, followed by an ice-cold methanol incubation. The final tissue fixation was performed with the Fixative reagent (Akoya Biosciences).

Stained coverslips were mounted onto the CODEX stage plate version 2 (Akoya) and secured onto the stage of a BZ-X810 inverted fluorescence microscope (Keyence). Reporter plates were prepared by adding fluorescently labeled oligonucleotides (Atto550, Cy5, AF750) made up in a reporter stock solution of nuclease free water, 10x CODEX buffer, assay reagent and nuclear stain to a black Corning 96 well plate (**Supplemental Table 1**). Automated image acquisition of tissue regions was performed at Enable Medicine using a CFI Plan Apo λ 20x/0.75 objective (Nikon) and fluidics exchange managed via the CODEX instrument and CODEX Instrument Manager software (CIM version 1.29.3.6, Akoya Biosciences), according to the manufacturer’s instructions, with slight modifications. Staining was evaluated for the expression of each marker in the panel. Non-specific staining was observed for EPCAM, SPP1 and CD163, which were excluded from downstream analysis resulting in a 25-plex panel.

### CODEX Image processing

Raw fluorescent TIFF image files were processed, deconvolved and background subtracted utilizing the CODEX Processor Software (Akoya Biosciences), and antibody staining was visually assessed for each biomarker and tissue region using the ImageJ software (Fiji, version 2.0.0). The TIFF hyper stacks were segmented based on DAPI nuclear stain, pixel intensities were quantified, and spatial fluorescence compensation was performed, which generated comma-separated value (**CSV**) and flow cytometry standard (**FCS**) files for downstream analysis.

### CODEX image registration

We excluded areas from neighboring normal liver, to ensure that CODEX analysis was performed on cells belonging to the mCRC TME. A pathologist evaluated images from hematoxylin and eosin (**H&E**) tissue sections, adjacent to the corresponding CODEX-stained sections. We used these annotated histopathology sections to distinguish normal liver parenchyma from tumor tissue regions. To determine which cells from the CODEX data were within normal liver tissue regions, nuclei were aligned between CODEX and H&E images. This was accomplished by aligning the CODEX nuclei segmentation images with the hematoxylin channel of corresponding H&E images using the image registration method described in HEMnet (26). Those cells within the annotated normal liver regions were excluded from further analysis.

### CODEX Context Assisted Cell Type Identification (CACTI)

A standard clustering procedure involves using cell type specific feature data to identify cell types. Due to the sparsity and noisy nature of measured CODEX data, expression of the index cell may not accurately represent the innate cell feature. By harnessing the information available about each cell’s local neighborhood to form a richer feature space, we improved the clustering of any specific index cell. We developed a method, Context Assisted Cell Type Identification (**CACTI**), which leverages spatial information during clustering.

For a given cell i, let X_i_ be its marker expression vector and Y_i_ be its spatial location. For a set of cells S, let X_S_ be the matrix that joins each vector *X_i_*_:*i*∈*S*_ row wise. Let Y_S_ be defined analogously. After normalization, the first step of CACTI is to identify the Delaunay neighbors of each index cell. These Delaunay neighbors will be a proxy for a cell’s local neighborhood. Letting Si be the set of Delaunay neighbors of index cell i, we define a niche feature to be a function f(X_Si_,Y_Si_). Some examples of f include Mean Expression, Distance weighted mean expression and Standard deviation. After calculating our niche features for each cell, we define our niche augmented data to be such that

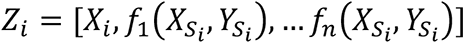

where *n* is the number of niche features calculated. One drawback of *Z* is that it might focus too much on the niche information compared to the underlying expression profile of the index cell. To overcome this, we introduce a weight parameter *λ* to the niche features and focus our analysis on

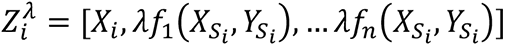

Let Zλ be the matrix that joins the individual 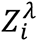 row wise. To determine *λ*, we perform low-resolution clustering. Individual Seurat objects were constructed from cell-feature matrices and spatial co-ordinates from each sample using the ‘Spatial’ Assay in Seurat. All objects were merged, and data was scaled using the ‘ScaleData’ function. Batch-correction was performed during clustering using the Harmony algorithm as outlined above. We used the first 10 principal components and a resolution of 0.2 for clustering. Let L be the low-resolution cluster assignment of X. Although CODEX data is noisy, we expect L to be a reasonable approximation of the major cell types present in our sample. For another cluster assignment of the same cells A such that |A| > |L|, we define E(A, L) to be the minimum classification error of A with respect to the low resolution clustering L. We recommend that A and L be generated by the same clustering algorithm (e.g. K-means, Louvain, etc.).

If our niche features contain perfect information pertaining to the true cell types, then given a clustering of Zλ, C(Zλ), we expect E(C(Zλ), L) to be small for all values of λ. On the other hand, if our niche features are independent of the true cell types, then E(C(Zλ), L) should be large for even moderate values of λ. Therefore, when choosing λ, we should choose one such that E(C(Zλ), L) is less than some level α. Mathematically, to find a suitable λ, we attempt to solve the following optimization problem:

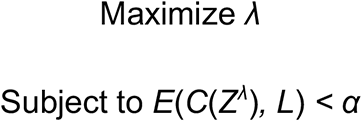

If *E*(*C*(*Z^λ^*)*, L*) is monotone in *λ* we can use a bisection algorithm to get the optimal *λ* within some desired degree of error.

Following low-resolution clustering and CACTI, clusters were annotated for cell types using lineage marker genes. We identified tumor epithelial cells (PanCK i.e., Pan Cytokeratin), CAFs (COL4A1, ACTA2), macrophages (CD68), endothelial cells (PECAM1), CD4 T cells (CD4), CD8 T cells (CD8) and Tregs (FOXP3). Cells co-expressing epithelial, or immune or stromal markers were filtered as artifacts (7.345% of total cells). A mixed cluster of macrophages and epithelial cells (11.81% of analyzed cells) was gated for macrophages expressing CD68>0 and PanCK <0.5 using the scaled data, with remainder cells classified as epithelial. A mixed cluster of epithelial cells, fibroblasts, and lymphocytes (2.92% of analyzed cells) was gated using scaled data for epithelial cells (PanCK > 0, CD45 <0) followed by lymphocytes (CD45 >0, COL4A1 <0) with the remainder cells classified as fibroblasts.

### Cellular proximity analysis

Let the total number of cells in our sample be *N*. Let *G* be a graph indexed by *N* nodes such that the weight of an edge between nodes *i* and *j* is the similarity between cells *i* and *j*. Now let *C*_1_, *C*_2_, and *C*_3_ be three sets of cells. The hypothesis test for proximity analysis can be formulated as:

H0 : *C*_1_ and *C*_2_ are on average equally similar to *C*_3._

HA : *C*_1_ is on average more similar to *C*_2_ than *C*_3_ is.

This hypothesis can be tested with a permutation test where we permute the labeling of members within sets *C*_1_ and *C*_2_ and calculate our test statistic

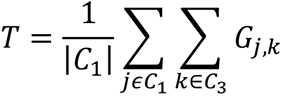

We reject the test when *T* is very large relative to the permutated test statistics. We wanted to examine whether fibroblasts are more spatially proximal to macrophages than other cells on average. This corresponds to letting *C*_1_ be the set of macrophages, *C*_3_ be the set of fibroblasts and *C*_(_ be all other cells. To model spatial proximity, we let the similarity between cells *i* and *j* be the Jaccard index between the K nearest neighbor sets of the two cells based on their spatial coordinates.

### Additional computational analysis

We used R packages tidyverse (v1.2.1), ggplot2 (v3.3.3), ggpubr (v0.40), broom (v0.5.2), viridis (0.5.1), pheatmap (v1.0.12), ComplexHeatmap (v2.9.3) (27), and stats (v4.0.5) in R v4.1.0 for additional analysis or visualization. Figures were additionally edited in Adobe Illustrator CS6 (v16.0.0).

## RESULTS

### Properties of the cellular TME of CRC metastases to the liver

Our study relied on three different approaches to characterize the CRC metastatic TME (**Fig. 1A**). We performed scRNA-seq analysis of mCRC tissue from surgical resections – these samples included patient matched normal liver and PBMCs (**Table 1**). The cohort consisted of 14 samples from seven patients. All tumors had adenocarcinoma histopathology. The only exception was P6198’s tumor which had mixed neuroendocrine adenocarcinoma (**MANEC**) histology. In addition, all tumors underwent clinical testing for microsatellite instability (**MSI**) via immunohistochemistry (**IHC**) for DNA mismatch repair proteins. All tumors were microsatellite stable (**MSS**). We examined the spatial organization of these cell states with CODEX on 15 MSS mCRCs found in the liver. Finally, we confirmed the presence of these cell states and interactions using an independent gene expression dataset of 93 mCRCs to the liver (3). With this RNA-seq data and a cell deconvolution method, we determined the association of cell states and specific clinical outcomes.

**Figure 1.**
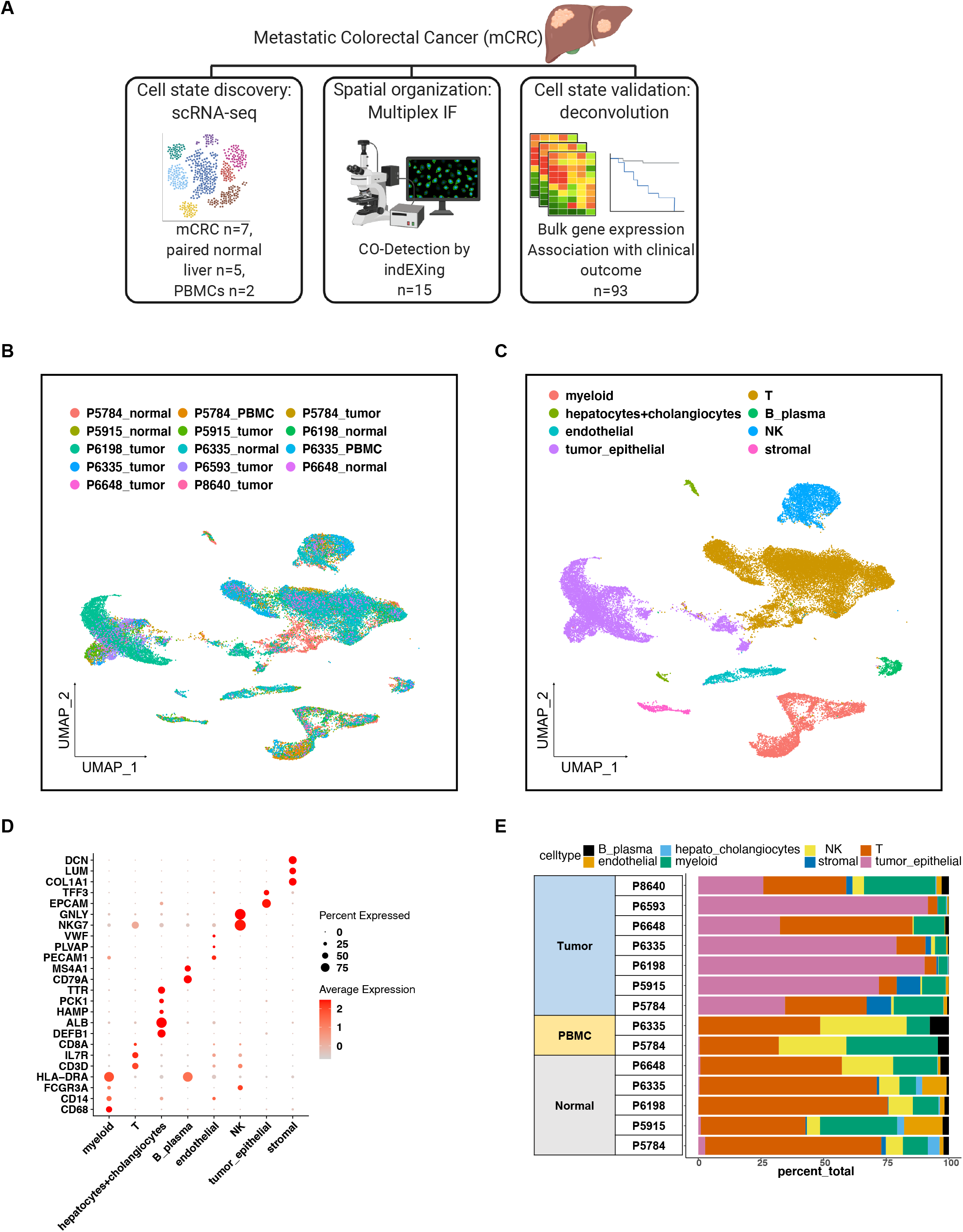
(A) Schematic representation of study design. (B-C) UMAP representation of dimensionally reduced data following batch-corrected graph-based clustering of all datasets colored by (B) samples and (C) cell type. (D) Dot plot depicting average expression levels of specific lineage-based marker genes together with the percentage of cells expressing the marker. (E) Proportion of cell types detected from each sample.

**Table 1.**
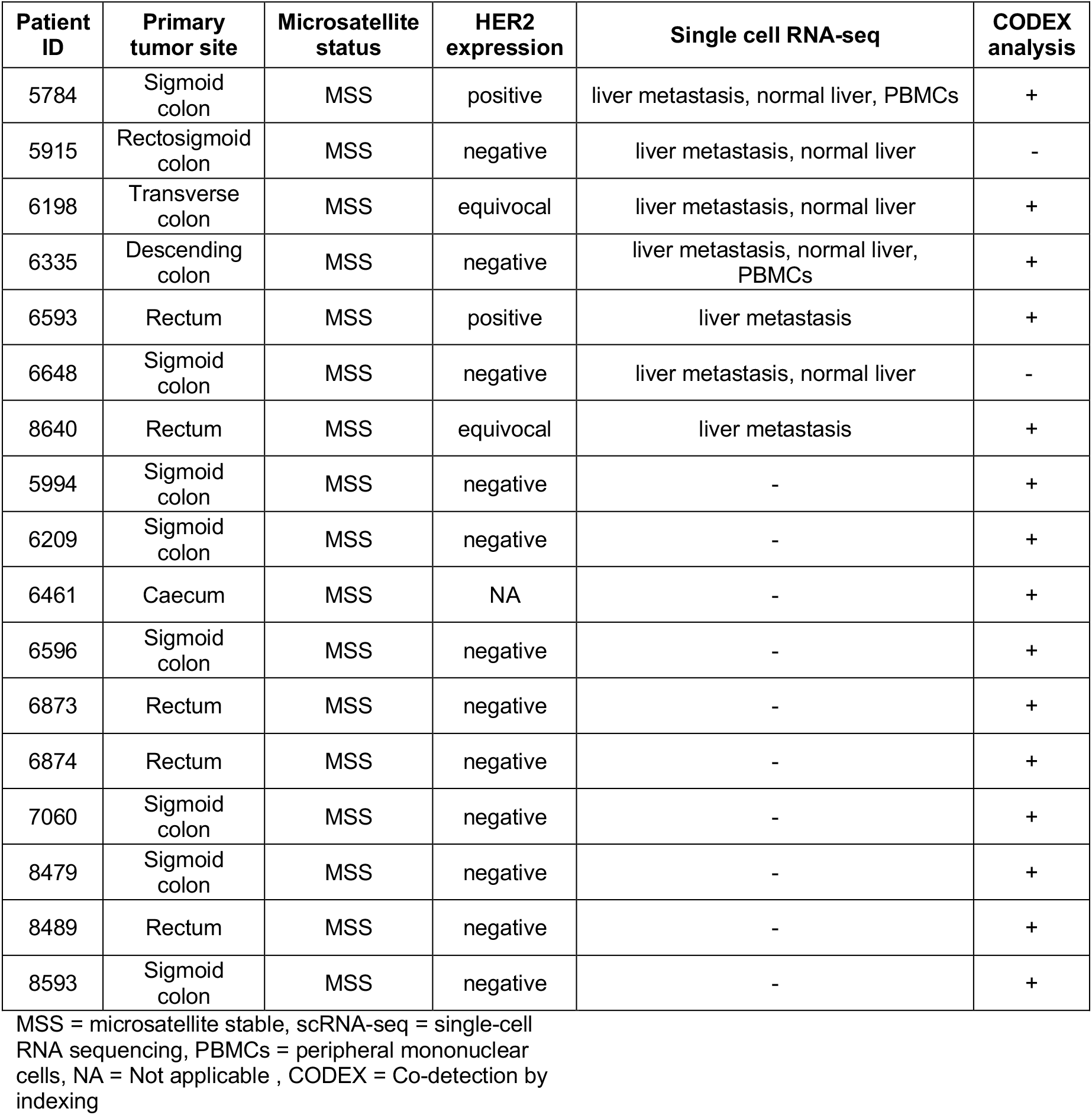
Metastatic colorectal cancers in the liver.

### Single-cell RNA analysis of CRC metastases in the liver

We sequenced a total of 44,522 single cells from these metastases. The data included 22,718 cells from normal liver, 14,848 cells from liver mCRCs, and 6,970 PBMCs (**Supplemental Table 2**). The total number of cells per sample ranged from 281 to 8,706 with the variation directly attributable to the size of the resected tissue sample. We filtered out poor quality data, eliminating cells with high mitochondrial genes indicative of cell death and computationally identified doublets (13, 28). This quality control step removed 12.2% of the total number of cells.

To identify the different cell types, we aggregated the data across all samples (**Methods**). We normalized the data, carried out steps to remove technical variation in sequencing depth and performed principal component analysis (11, 12). Data sets from different experimental batches were integrated with a k-means clustering method implemented in the Harmony program (14).

We used Uniform Manifold Approximation and Projection (**UMAP**) (29) to visualize the resulting clusters. Most cell clusters had contributions from different samples (**Fig. 1B**), indicating that there were no obvious batch effects during clustering. We confirmed this computationally by examining a similarity metric called the Adjusted Rand Index (**ARI**) (30). Comparison of cluster assignments with experimental batch had an ARI of 0.06, a low value indicating near-random assignment. In summary, we confirmed that cluster assignments were not the result of experimental batch effects.

Clusters were annotated with major cell types according to the expression of established marker genes for specific cell types (17, 31, 32) (**Fig. 1C, 1D**). From this data, we identified normal hepatocytes (*ALB*, *HAMP*, *PCK1*, *TTR*), cholangiocytes (*DEFB1*), tumor epithelial cells (*TFF3, EPCAM*), endothelial cells (*VWF*, *PLVAP*, *PECAM1*) and fibroblasts (*DCN*, *LUM*, *COL1A1*). Representing the immune cell types, we detected myeloid lineage cells (*CD14, FCGR3A, CD68, HLA-DRA*) that included macrophages and dendritic cells, T lymphocytes (*CD3D*, *IL7R*, *CD8A, NKG7*), NK cells (*GNLY*, *NKG7*) and B cells (*CD79A*, *MS4A1*).

Depending on the size of the tissue sample, the absolute number of cells, their types and their proportions varied (**Fig. 1E**, **Supplemental Table 3**). Subsequently, we performed secondary clustering analysis with batch-correction for each cell lineage to determine their gene expression properties and extrapolate more granular details about their cell state.

### Gene expression properties of metastatic tumor epithelium

The CRC epithelial cells formed patient-specific clusters among the different mCRCs, reflecting the genomic diversity of these cancers (**Supplemental Fig. 1A**). We determined differential gene expression among mCRCs from the different patients. Each tumor had its own set of differentially expressed genes including *FABP1*, *OLFM4*, *KRT20*, *CEACAM5* and *CEACAM6*: these genes have been previously associated with CRC (31) (**Supplemental Fig. 1B**). We also detected high expression levels of *TSPAN8* and *HES1* which are indicators of a cancer-related stem cell state and properties of invasion (33, 34). Elevated *ERBB2* expression was detected in P5784’s and P6593’s tumor epithelial cells - this result was corroborated by IHC results that also confirmed ERBB2 overexpression (**Table 1**). In addition to marker genes associated with colorectal adenocarcinoma, P6198’s MANEC metastasis had significantly increased expression of *DEFA5* and *DEFA6* genes. The high expression of these genes occurs in small intestinal neuroendocrine tumors (35).

### Tumor epithelial cell aneuploidy and chromosomal imbalances

We evaluated the extent of chromosomal scale copy number variations (**CNVs**) among the tumor epithelial cells. Large copy number alterations that extend up to entire chromosome arms are also referred to as allelic imbalances. This analysis relied on the InferCNV program which processes each cell’s gene expression across a given chromosome, compares the results with reference diploid cells and provides somatic CNV changes (21).

The tumor epithelial cells in all mCRCs had significant levels of chromosome scale CNVs and allelic imbalances extending across the chromosome p or q arms (**Supplemental Fig. 1C**). These large-scale chromosomal events are indicators of aneuploidy and have been associated with mCRC (36). There was no discernible copy number variation from the other normal cell types. This result confirmed the identity of the cancer epithelial cells and indicated that the mCRCs belonged to the molecular subtype associated with chromosomal instability (**CIN**) (37). Notably, all tumors had undergone IHC for DNA mismatch repair proteins and were confirmed to be MSS, which is consistent with these mCRCs being CIN.

Citing some frequent copy number alterations, we observed chromosome allelic imbalance across chromosome arm 7p across all tumors. A deletion involving the chromosome 8p arm was observed among five out of seven mCRCs. There was a series of other frequent allelic imbalances involving CNV gains. Two tumors (P6648, P6335) had amplifications in chromosome 13. Three tumors (P6648, P6593, P6198) had chromosome 19 allelic imbalances. Four tumors (P6648, P6335, P5915, P5784) had imbalances in chromosome 20. All these chromosomal alterations have previously been identified as markers for increased risk of metastasis in CRC (38). The mCRC (P6198) with MANEC histopathology had genomic instability events including loss of chromosome 8 that is a frequent event among colorectal adenocarcinomas (39).

### Myeloid lineages in mCRC, normal liver and PBMCs

We examined the myeloid cell populations among the different samples following a secondary clustering analysis (**Fig. 2A, B**). The myeloid cell populations had clusters associated with the tissue source. The matched liver tissue had normal myeloid cells present in multiple distinct clusters. The matched peripheral blood had normal monocytes that cluster separately without overlap from other macrophage types. The macrophages from the mCRC samples distinctly separated from the macrophages in the matched normal liver tissues and peripheral monocytes. Specifically, mCRCs macrophages were represented among Clusters 1 and 3 (**Fig. 2A, B**).

**Figure 2.**
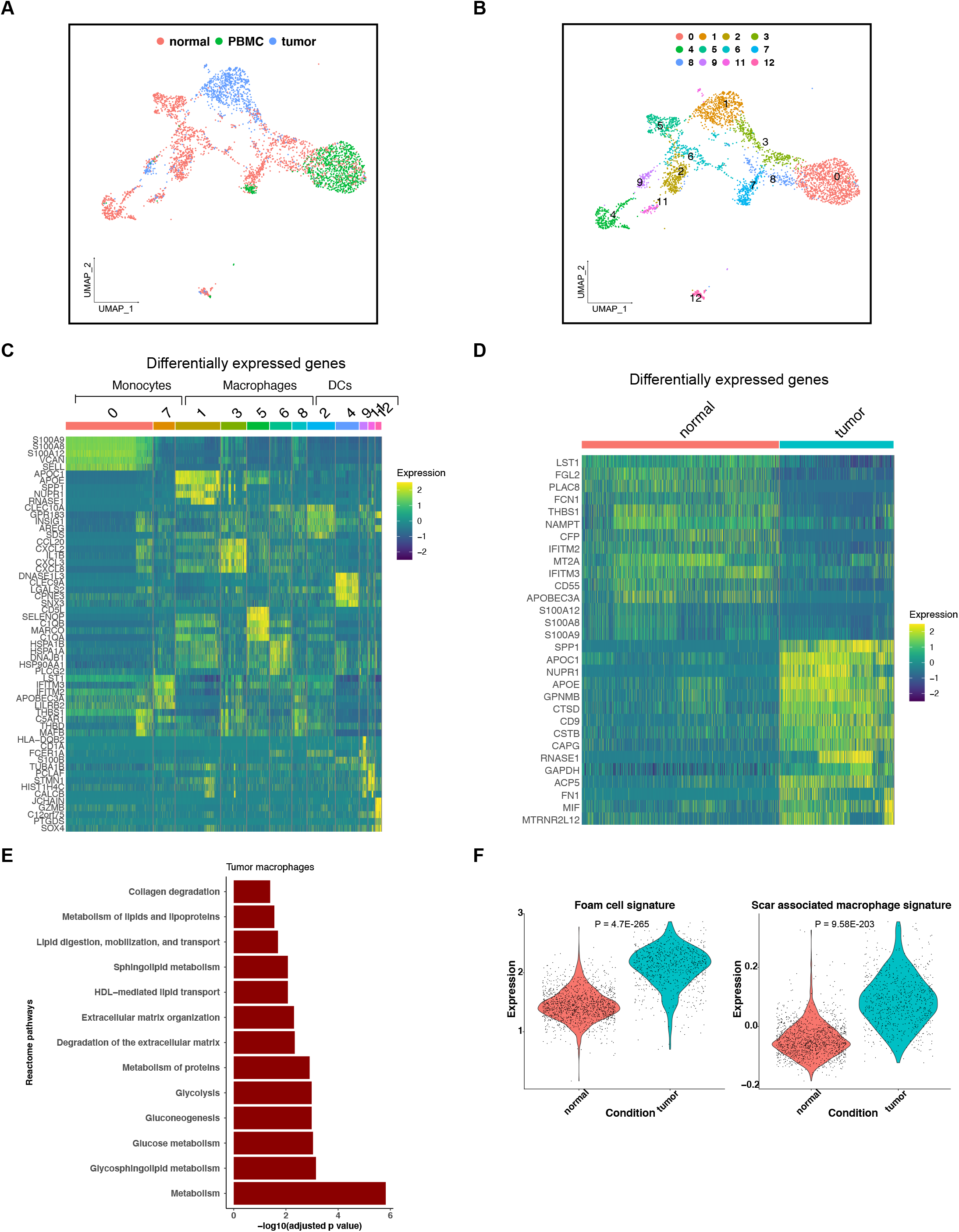
(A-B) UMAP representation of dimensionally reduced data following batch-corrected graph-based clustering of all myeloid lineage cells annotated by (A) condition and (B) cluster numbers. (C) Heatmap depicting expression of five highest significantly expressed genes (adjusted p-value < 0.05) per cluster. (D) Heatmap depicting the expression of highest top 15 significantly expressed genes in normal and tumor macrophages (adjusted p-value < 0.05). (E) Selected differentially enriched reactome pathways in tumor macrophages. (F) Violin plots depicting the expression of gene signatures of foam cells or scar associated macrophages in normal and tumor macrophages with T-test p-value.

Next, we determined which genes defined the specific myeloid clusters. The PBMC monocytes expressing either *CD14* or *FCGR3A* (CD16) highly expressed S100A family genes (**Figure 2C**). Dendritic cells expressed the HLA genes, *CD1C*, *CLEC9A* and *IDO1* among others. Intrahepatic macrophages included normal Kupffer cells that expressed *CD5L*, *MARCO*, *LIPA*, *MAF*, *VCAM1* etc. (17, 40).

There was a population of tumor-associated macrophages with high expression levels of *SPP1*. These macrophages were present in Clusters 1 and 3. The macrophages in Cluster 1 also expressed *APOC1*, *APOE, RNASE1* and others. The macrophages in Cluster 3 expressed chemokines genes such as *CXCL8*, *IL1B*, *CCL20* and others. The elevated expression of *SPP1* has been identified among tumor-associated macrophages across several cancers including primary CRC (41). *SPP1* encodes for an integrin binding glyco-phosphoprotein. *SPP1* overexpression is observed in cancer and is associated with a poor prognosis (42). We refer to this specific cell type as *SPP1+* tumor associated macrophages.

### Reprogrammed tumor associated macrophages have inflammatory fibrosis and lipid metabolism features

Macrophages display a high degree of plasticity, which is related to assuming different functional properties. These changes in the cell states are generally referred to as reprogramming. Our analysis discovered that *SPP1*+ tumor associated macrophages had gene expression signatures reflecting two reprogrammed functional states: 1) scar associated macrophages present in fibrotic cirrhotic livers; 2) foamy macrophages that have engulfed high levels of low-density lipoprotein.

We compared the gene expression signature of the metastatic TME macrophages to the other macrophage types present in normal liver tissue (**Fig. 2D**, **Supplemental Table 4**). The *SPP1*+ tumor-associated macrophages had elevated expression levels of *APOC1*, *APOE*, *TREM2*, *FN1, LGALS3, FTL, CD9, CTSB,* etc. (p value < e-72). These genes are notable for defining specific cell properties. Namely, macrophages with increased *SPP1, TREM2*, *FN1* and *LGALS3* expression occur in fibrotic diseases such as pulmonary fibrosis and cirrhosis (43, 44). From studies of primary CRCs in the colon, macrophages expressing *SPP1* and *CTSB* were associated with construction of a collagenous ECM (45). *LGALS3* encodes for a member of the galectin family of carbohydrate binding proteins. It plays a role in macrophage polarization and fibrosis in inflammatory diseases (46). *TREM2* functions as a molecular regulator of the foam cell phenotype in macrophages (47). Elevated TREM2 expression was also identified in macrophages in liver cirrhosis, indicating a pro-fibrotic function (44). Meanwhile, high expression of *APOE* and *APOC1*, encoding for lipoproteins, indicated higher levels of cholesterol metabolism. Similar expression features are observed in foam cell macrophages located in atherosclerotic plaques, an obstructive lesion of arterial vessels (18). In summary, TME macrophages in the liver had gene expression signatures observed in inflammatory fibrosis and lipid metabolism.

We applied different expression analysis methods to confirm the functional states of these reprogrammed TME macrophages. Using the enrichR program (16), we performed a pathway analysis on the differentially expressed genes in TME macrophages to identify the biologically relevant processes regulated by them. We detected significant enrichment of terms relating to both extracellular matrix (**ECM**) organization and metabolism. Different metabolic pathways were enriched including glycosphingolipid metabolism, glucose metabolism and HDL-mediated lipid transport (**Fig. 2E**, **Supplemental Table 5**). Next, we quantified the expression signature from foamy macrophages (18) and cirrhotic scar associated macrophages (44) (**Supplemental Table 6**). Compared to normal hepatic macrophages, mCRC macrophages had significant enrichment of both these gene signatures (**Fig. 2F**). These results overlap with the results of the differential gene expression analysis.

As an alternative approach for evaluating the macrophage properties, we used the EcoTyper program. It employs a non-negative matrix factorization (**NMF**) on gene expression data such as scRNA-seq to identify cell states (24). EcoTyper does not rely on single cell clustering. The NMF analysis of the mCRC data identified a distinct expression signature enriched for tumor macrophages (**Supplemental Fig. 2**). This tumor macrophage signature included *SPP1*, *GPNMB*, *APOC1*, *APOE*, *TREM2*, *CTSB*, *LGALS3*, *FTL*, etc. These genes overlapped with the results from the differential expression between tumor-associated and normal macrophages (**Fig. 2B**, **Supplemental Table 4**). Overall, this result confirmed the identification of a distinct macrophage cell state in the mCRC TME with fibrogenic properties and altered metabolism.

### Stromal cell components in the metastatic microenvironment in the liver

We characterized the different stromal cells present in the mCRC microenvironment in the liver using reclustering with batch correction. The clustering analysis showed that the stromal cells from the mCRCs separated from those in the normal liver (**Fig. 3A**). Among the different clusters there were three major cell types which included fibroblasts, endothelium and hepatic stellate cells (HSCs) (**Fig. 3B**).

**Figure 3.**
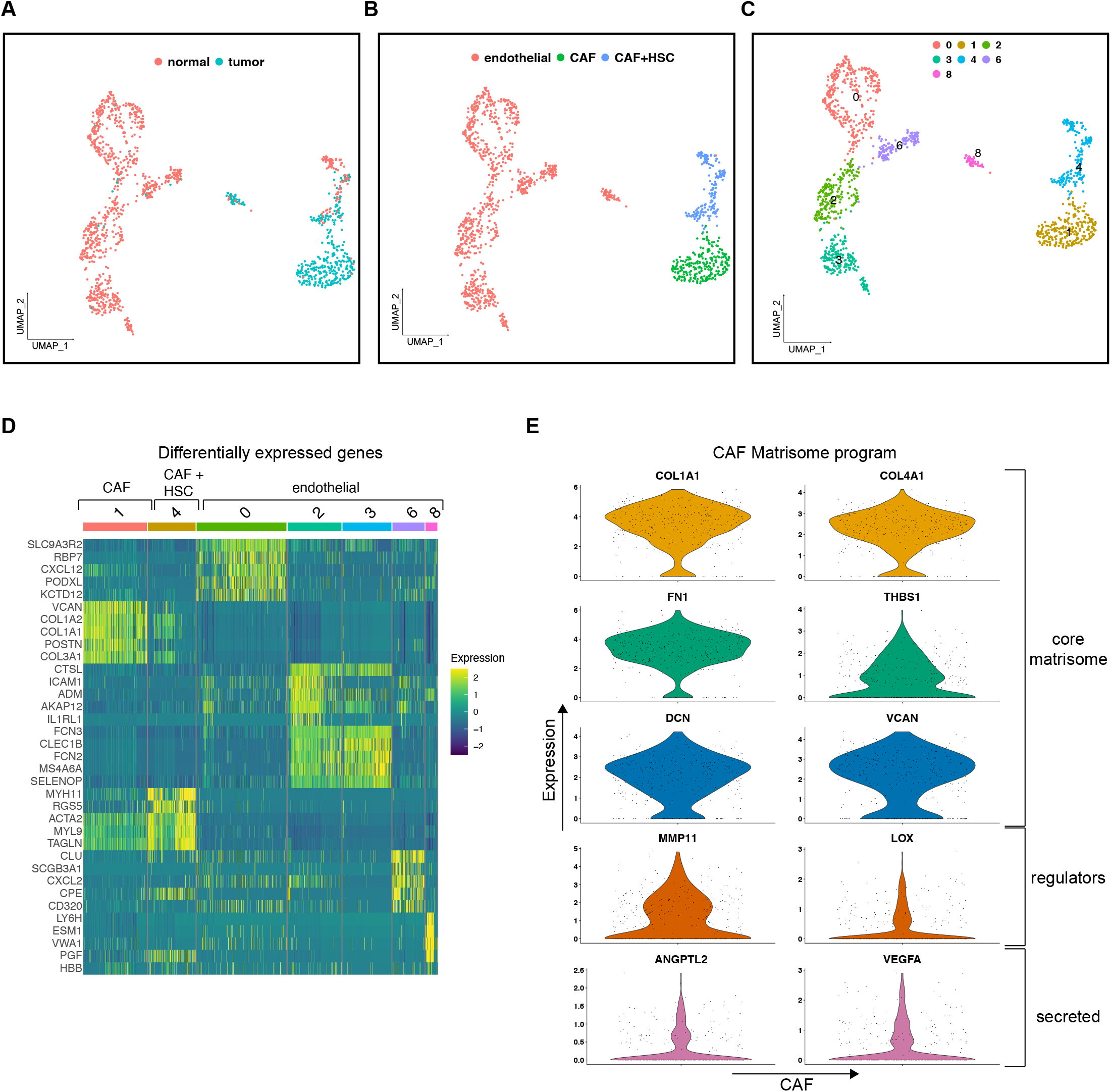
(A-C) UMAP representation of dimensionally reduced data following batch-corrected graph-based clustering of all stromal lineage cells annotated by (A) condition, (B) cell types and (C) cluster numbers. (D) Heatmap depicting expression of five highest significantly expressed genes (adjusted p-value < 0.05) per stromal cell cluster. (E) Violin plots depicting the expression of selected matrisome components in differentially expressed genes in CAFs.

Fibroblasts associated with mCRC were present in Clusters 1 and 4 (**Fig. 3B, 3C**). Cluster 1 only contained fibroblasts from mCRC and was distinctly separated from the fibroblasts of normal hepatic tissue. These cells were characterized by elevated expression of ECM-related genes such as those involved in collagen synthesis, *POSTN*, *FN1*, *MGP*, etc. (**Fig. 3C**). Therefore, Cluster 1 had the attributes of cancer-associated fibroblasts (**CAF**).

The fibroblasts in Cluster 4 had high expression of *ACTA2* (Cluster 4). These cells overlapped with HSCs from normal liver. Additional genes with differential expression included *TAGLN*, *MYL9* and *IGFBP7* – these genes are expressed in activated HSCs (17). HSCs are quiescent fibroblasts, occupying a specific cellular niche in the liver. Inflammatory processes activate these stellate cells, allowing them to proliferate and migrate. Our analysis also identified endothelial cell subsets (Clusters 0, 2, 3, 6 and 8) which were present in normal liver.

### Fibroblasts in the metastatic TME have a tumor-promoting ECM expression signature

Next, we compared the set of CAF differentially expressed genes with the components of the ‘matrisome’, a term that refers to the core ECM components. The matrisome includes fibronectins, collagens, laminins, proteoglycans, etc. which are associated with the ECM structure and its secretion (**Fig. 3E**, **Supplemental Table 7**) (48). The CAFs had a matrisome program based on the differential expression of several collagen genes (*COL1A1*, *COL3A1*, *COL5A1*, etc.), a variety of ECM glycoproteins (*FN1*, *POSTN*, *SPARC*, *THBS1*, etc.) and proteoglycans (*BGN*, *VCAN*, etc.). These cells also highly expressed ECM regulator genes including *MMP11*, *MMP14*, *TIMP1*, *LOXL1* and *LOXL2*. These genes are involved in ECM remodeling. The ECM composition influences physical properties such as stiffness and contributes to tumor growth and drug resistance (49).

The CAFs were also denoted by the expression of secreted growth factors including *VEGFA*, *PDGFA* and *PDGFC*. These genes promote tumor growth and enable immune evasion (50). For example, *VEGFA* is involved in supporting the migration of cancer cells and facilitates metastasis.

### The metastatic TME has an immunosuppressed T cell milieu

For all mCRCs, there was a lack of tumor-reactive CD8 T cells in the TME of the liver. Moreover, we detected regulatory T cells (**Tregs**) in the TME. These cell features are the hallmarks of an ineffective anti-tumor response. We analyzed all lymphocytes from mCRCs, PBMCs and normal tissue. Based on marker gene expression, we detected CD8 T cells, CD4 T cells, NK/NK-like cells (gamma delta T, NK-T, MAIT atypical, Tregs, plasma and B cells (**Supplemental Fig. 3A, 3B**). The T and NK cells in PBMCs clustered separately from the same cell types in the liver, indicative of tissue-specific transcriptional differences, which we have also observed in gastric tissue (**Supplemental Fig. 3C**).

CD8 T cells from tumors co-clustered with those from normal liver, indicating their gene expression signatures were similar. These cells expressed markers of previously described tissue resident cells in the liver (17) including *GZMK*, *CCL5*, *CCL4L2* and *CD69*. Notably absent were CD8 T cells with features of tumor-reactivity such as expression of *ITGAE*, *ENTPD1* and *CXCL13* (**Supplemental Fig. 3B**) (51). We confirmed the transcriptional similarity between tumor and normal liver CD8 T cells using the NMF-based, non-clustering EcoTyper algorithm previously described (**Supplemental Fig. 3D**). Cell states of CD8 T cells in the TME overlapped with those of normal liver CD8 T cells. This result supports the conclusion that CD8 T cells in the TME are quiescent bystanders. We evaluated the expression of a cytotoxic gene signature among these CD8 T cells. The cytotoxicity signature among the tumor CD8 T cells was significantly lower than those in normal liver CD8 T cells (P = 1.23E-11, **Supplemental Fig. 3E**). In summary, for all the mCRCs, the results from the CD8 T cells and the presence of Tregs indicated an immunosuppressed TME lacking anti-tumor activity.

### TME fibroblasts and macrophages influence the T cells in the mCRC

Using the single cell RNA-seq data, we characterized the receptor-ligand networks present in the liver TME. We discovered intercellular interactions among non-immune and immune cell types that facilitate T cell exclusion and exhaustion. For this analysis, we used the program CellChat to identify cell-type specific receptor-ligand interactions and construct a mCRC TME interactome (22). This algorithm identities the differentially over-expressed ligand genes and their complementary receptors for each cell type, quantifies each interaction with a probability value and delineates the significant interactions by randomly permuting the cell type labels.

Macrophages and CAFs were noted to have expression of specific ligands that contribute to T cell exclusion and exhaustion. Specific interactions identified within these pathways included the fibroblast-lymphocyte *CXCL12*-*CXCR4* receptor ligand pair (**Supplemental Fig. 4A**). CXCL12 ligand and its co-receptor CXCR4 regulate the mobilization of immune cells into tissues (52). We also identified expression of *NECTIN2* from fibroblasts and endothelial cells. This ligand binds to immune checkpoint TIGIT present in T cells.

As described previously, we found that the TME macrophages expressed *SPP1* – this ligand suppresses T cell activation via interaction with *CD44* (53). This ligand also interacts with the integrin receptor family, thus cross networking with CAFs (54). Macrophages expressed the *CD86* ligand that maintains the regulatory phenotype and survival of Tregs via interaction with *CTLA4* (55). CAFs and macrophages expressed *VEGFA* and *VEGFB* that can mediate angiogenesis (56).

We visualized these interactions as lines between different cell types, with their width scaled by the number of interactions mediated by the sender cell. CAFs were the most prolific communicators in the TME, dominating the top 10% of all cell-to-cell interactions (**Supplemental Fig. 4B**).

### Networking of macrophages and CAFs mutually influence their cell states

We discovered that macrophages and CAFs affect each other’s gene expression programs via specific receptor-ligand interactions. Our analysis used the NicheNet algorithm that can predict ligands from a sender cell type that regulate target gene expression in a receiver cell type via ligand-receptor interactions (23). This analysis can thus identify intercellular communication that influences the transcriptional phenotype of a target cell. We visualized these interactions as a Circos plot with ligands from sender cells that affect downstream target gene expression in a receiver cell.

We first identified ligands from cells in the TME that can result in expression of matrisome genes in mCRC CAFs (**Fig. 4A**). One of the highest ranked genes was the established ECM regulator gene *TGFB1*, which was derived from NK cells, validating this approach (57). Several ligands were derived from macrophages including *SPP1*, *IL1B*, *TNF*, *MMP9* and *CCL2*. These ligands have the potential to regulate target gene expression of several core matrisome genes including the collagen family. Other CAF matrisome target genes for macrophage ligands included *MMP2* and *VEGFA*. This result further supports our finding that the reprogrammed *SPP1*+ macrophage cell state promotes fibrosis in the mCRC TME. Additionally, several ligands were expressed by CAFs themselves, indicating autocrine signaling. These ligands included *AGT*, *TGFB3*, *CTGF*, *CCL2*, *FGF1*, *HGF*, *CXCL12* and *CSF1*.

**Figure 4.**
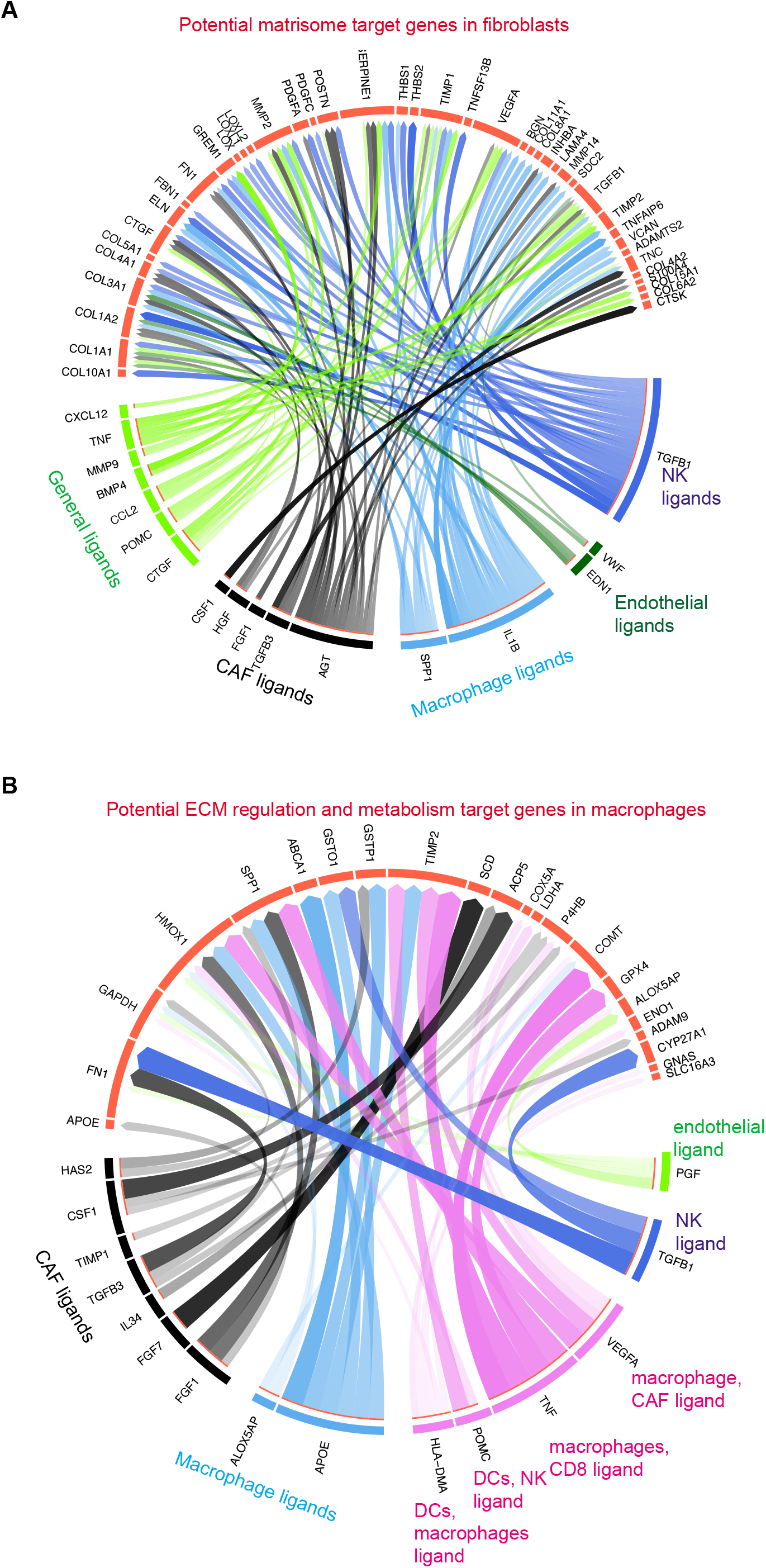
(A-B) Predicted ligands that regulate respective target genes in (A) CAFs and (B) macrophages. Ligands are annotated by the cell type that expresses them. General ligands indicate ligands expressed by more than one cell type. Edges are scaled by the inferred regulatory potential of the interaction.

Next, we examined which ligands can lead to the reprogrammed macrophage cell state with features indicative of inflammatory fibrosis and lipid metabolism. The top ranked ligands included *FGF1*, *CSF1*, *PGF*, *TGFB3* and *TIMP1*; all were derived from CAFs. (**Fig. 4B**). These ligands can target macrophages and regulate the expression of *SPP1*, *FN1* and *APOE*. Hence, ligands from CAFs have the potential to reprogram the mCRC macrophages via ligand-receptor interactions. Overall, these results pointed towards the presence of a signaling network between TME macrophages and CAFs. This intercellular communication influences the transcriptional phenotype of both cell types.

### Spatial characterization of the mCRC TME in the liver

To determine the spatial cellular characteristics of the mCRCs, we used CODEX multiplexed imaging. This approach uses antibody multiplexing on tissue sections, enabling cell type identification at a single-cell resolution (**Fig. 5A**) (58). This spatial imaging allowed us to ask specific questions about cell types and cellular proximity in the liver mCRC TME.

**Figure 5.**
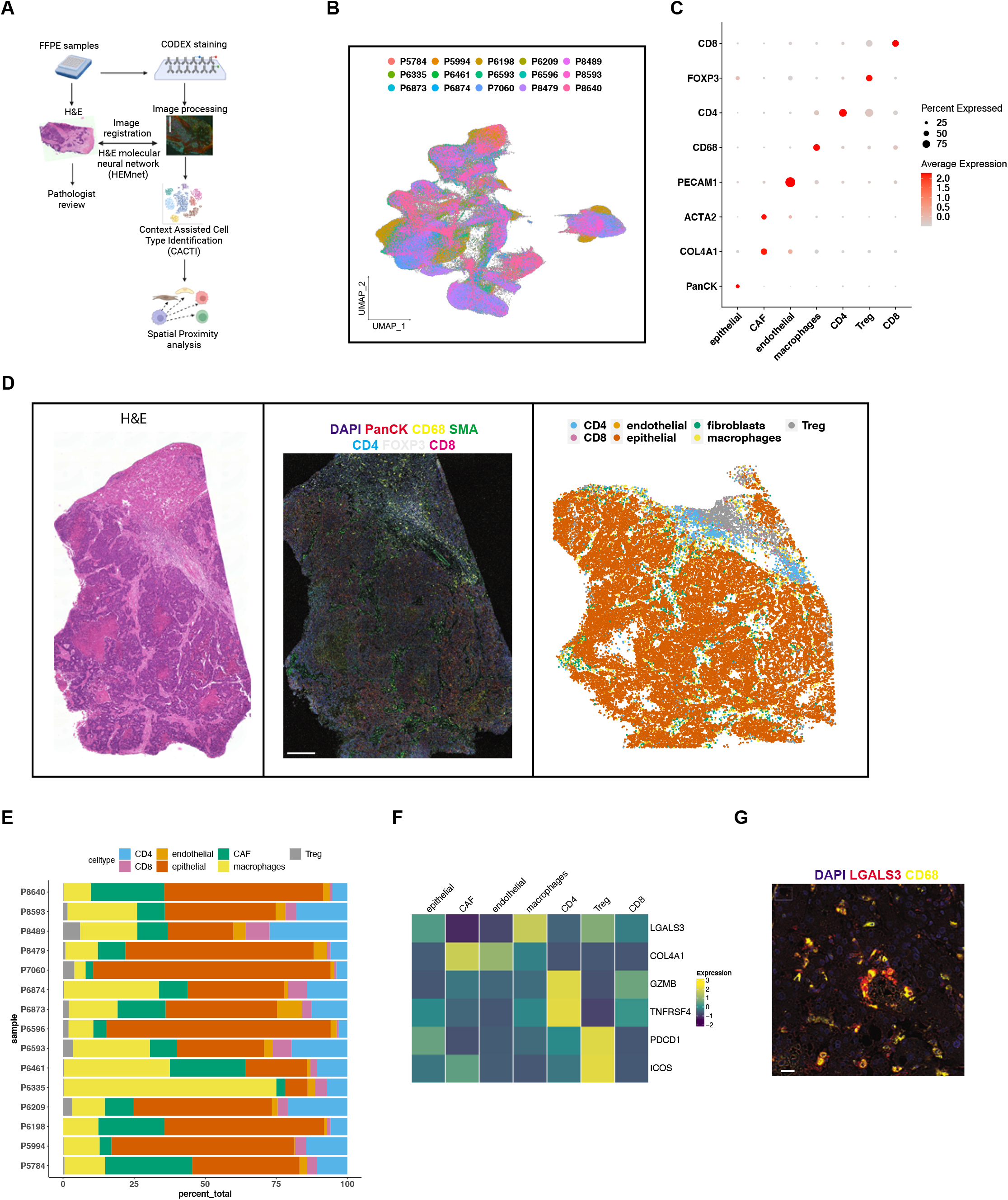
(A) Schematic representation of CODEX analysis. (B) UMAP representation of dimensionally reduced CODEX data following batch-corrected graph-based clustering of all datasets colored by samples. (C) Dot plot depicting average expression levels of specific lineage-based marker proteins together with the percentage of cells expressing the marker. (D) Example of P7060 tumor with adjacent H&E section (left panel), CODEX staining of selected cell lineage markers (middle panel) and graphical representation of identified cell types in image data (right panel). Scale bar = 1.07 mm. (E) Proportion of cell types detected from each sample. (F) Heatmap depicting average expression of selected proteins across all samples in respective cell types. (G) Example of P7060 tumor with CODEX staining of selected markers. Scale bar = 90 μm.

We used a 25-plex antibody panel (**Supplemental Table 1**). This panel included lineage markers to identify specific cell types including tumor epithelial cells (PanCK i.e. Pan Cytokeratin), CAFs (COL4A1, ACTA2), macrophages (CD68), endothelial cells (PECAM1), CD4 T cells (CD4), CD8 T cells (CD8) and Tregs (FOXP3). A subset of antibodies were specific for proteins expressed in different cell states identified in our scRNA-seq analysis. These antibodies included one which recognizes LGALS3, a marker of the inflammatory fibrosis phenotype seen in the *SPP1+* tumor associated macrophages. We also examined the expression of immune checkpoints PDCD1 (PD-1) and ICOS; and co-stimulatory molecule TNFRSF4 (OX40) that characterize dysfunctional CD8 T cells as well as Tregs. Further, we examined the expression of the cytotoxic effector molecule GZMB.

We analyzed both the CODEX and hematoxylin and eosin (**H&E**) images for each tumor. The mCRC tissues underwent pathology review from an adjacent H&E section. The annotation outlined the boundaries between the tumor and adjacent normal liver parenchyma. Next, we processed the CODEX image data to excluded adjacent normal liver. For this step, the analysis used the HEMnet program which processes both the H&E and CODEX images to map the nuclei from CODEX to H&E images with corresponding pathological annotation (26). This enabled the exclusion of images covering the normal liver parenchyma from downstream analysis. Hence, our analysis could be restricted to tumor cells and the surrounding TME.

After image processing, there were a total of 330,893 single cells from 15 mCRCs (**Table 1**). We first clustered these cells using low resolution batch corrected clustering implemented in the Harmony algorithm (14). Cell clusters had contributions from different tumors. This result indicates an adequate removal of batch effects (**Fig. 5B**). Due to the sparsity and noisy nature of measured CODEX data, feature expression may be inadequate to resolve cell types based on clustering. We leveraged the spatial information of each index cell during the clustering process. This method, Context Assisted Cell Type Identification (**CACTI**) (**Methods**), improved the cell assignments per cluster following batch corrected clustering.

Based on the antibody staining patterns, we identified tumor epithelial cells, CAFs, macrophages, endothelial cells, CD4 T cells, CD8 T cells and Tregs. We verified cell type assignments by comparing corresponding H&E images (**Fig. 5C, D, Supplemental Fig. 5**). The different cell types had varying proportions across the mCRCs (**Fig. 5E**). Five samples had both scRNA-seq and CODEX results from different parts of the tumor. Proportions of cell lineages identified in these samples using the two methods demonstrated a moderate correlation (Pearson correlation coefficient 0.39, p = 0.02) (**Supplemental Fig. 6A)**.

### Validation of cell states in the mCRC TME

Having identified cell types, we examined the expression of specific markers characterizing distinct cell states. These markers were identified from the scRNA-seq results. TME macrophages had high expression of LGALS3, compared to other cell types (**Fig. 5F, G**). Macrophages across all patients had high correlation (Pearson correlation coefficient 0.76, p = 0.00099) between the expression of LGALS3 and lineage marker CD68 (**Supplemental Fig. 6B**). This result independently confirmed the *LGALS3* high *SPP1+* macrophages we identified in the scRNA-seq data.

CAFs had high expression of COL4A1, compared to other cell types (**Fig. 5F**). High co-expression of COL4A1 and ACTA2 was noted across CAFs from all patients (Pearson correlation coefficient 0.76, p = 0.001) (**Supplemental Fig. 6C**). This result supports the identification of the matrisome program identified in our scRNA-seq analysis of CAFs.

Among the lymphocytes, CD4 T cells had high protein expression levels of GZMB and TNFRSF4 (OX40). Tregs highly expressed immune checkpoints PDCD1 (PD-1) and ICOS. The CD8 T cells did not express these markers. These results support our finding that the mCRC TME lacks anti-tumor dysfunctional CD8 T cells expressing cytotoxic effectors, checkpoints, or costimulatory molecules. We detected Tregs in all samples (**Fig. 5E, Supplemental Fig. 5**). This result supports the immunosuppressed T cell milieu of the mCRCs observed in the single cell analysis.

### Spatial proximity between macrophages and fibroblasts in the metastatic TME

In our scRNA-seq analysis, we determined that *SPP1*+ tumor associated macrophages had a fibrogenic gene expression program. Moreover, we identified intercellular communication between macrophages and CAFs. We hypothesized that these two cell types are in physical proximity in the local cellular neighborhood of the TME. This proximity would facilitate any paracrine interactions. We examined the spatial proximity between macrophages and CAFs in the CODEX dataset.

We used this analysis on twelve samples which had the highest tissue integrity and minimal areas of necrosis (**Supplemental Fig. 5**). The latter feature lowers the quality of the image and acts as a confounding factor for the analysis. For each sample, we tested the hypothesis if CAFs were more spatially proximal to macrophages than any other group of cells on average. To test this hypothesis, we used a permutation test to permute cell labels from all macrophages, lymphocytes, epithelial and endothelial cells. We then examined if CAFs and each cell label was a mutual nearest neighbor based on their spatial co-ordinates. Hence, we could test if CAFs were significantly closer to macrophages than any other cell.

We detected significant spatial proximity between CAFs and macrophages in nine mCRC samples, compared to proximity between CAFs and all other cell types (permutation test p < 2.2E-16) (**Supplemental Table 8**). Hence, macrophages and CAFs are located spatially close to one another in the mCRC TME. This can enable paracrine interactions that influence their cell states. Overall, this result provides additional support for our scRNA-seq analysis that identified intercellular communication between macrophages and CAFs.

### Impact of CAFs on clinical outcomes in an independent mCRC dataset

To validate our findings from our single cell discoveries, we analyzed gene expression data from 93 mCRCs resected from the liver (3). These tumors had undergone conventional RNA sequencing (**RNA-seq**). Notably, 96.6% of these tumors were MSS. Thus, the tumors had the same liver-based TME features as the cohort used for scRNA-seq.

We utilized a deconvolution method, CIBERSORTx, to infer cell lineage fractions in this dataset (25). Using this method, one can generate cellular fractions in a bulk gene expression dataset using single-cell profiles. We generated a gene signature matrix per cell lineage derived from cells specific to tumor samples, while excluding normal liver tissue and PBMCs (**Fig. 6A**). This analysis included tumor epithelial cells and TME-specific CAFs, *SPP1*+ macrophages, DCs, endothelial cells, CD8 T, CD4 T, Treg, NK, B and plasma cells. Applying this signature matrix to bulk gene expression datasets resulted in quantification of cellular fractions of each lineage per sample. We successfully obtained cellular fractions for all lineages (deconvolution p < 2.2E-16 with 1000 permutations). Hence, tumor-specific single-cell signatures could successfully be deconvoluted in an independent mCRC gene expression dataset.

**Figure 6.**
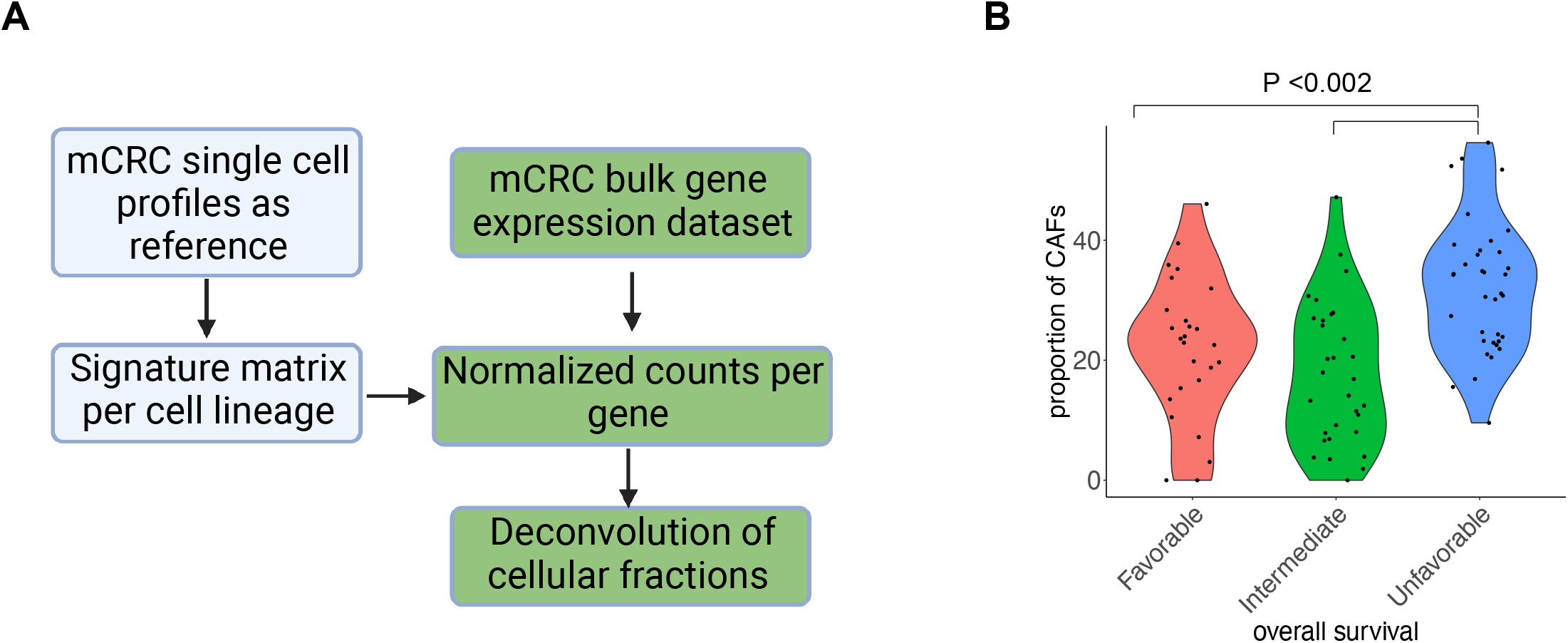
(A) Schematic representation of deconvolution of cellular fractions from external bulk RNA-seq dataset. (B) Violin plot depicting abundance of CAFs per patient with patients grouped according to overall survival sub-group. Comparisons were made by ANOVA with Tukey HSD.

We also assessed the impact of CAF abundance on prognosis. This external dataset from liver mCRCs identified three sub-groups with favorable, intermediate, and unfavorable overall survival (**OS**) (3). The sub-group with unfavorable OS had significantly higher proportion of CAFs (ANOVA FDR p < 0.002) (**Fig. 6B**). Hence, a TME phenotype characterized by high number of CAFs is associated with a poorer clinical outcome.

## DISCUSSION

Our study revealed novel networking between macrophages and fibroblasts in the mCRC TME. Using scRNA-seq we identified distinct communication programs between these cells with the potential to mutually influence their cell states. The potential for macrophage-fibroblast interactions is further supported by spatial analysis using multiplexed immunofluorescence that showed significant proximity between these cells. This interactome has the potential for modulating the metastatic niche, which represents the “soil” component of the metastatic cascade that allows tumor seeding.

We determined that TME macrophages had a gene expression signature that included *SPP1*, *APOE*, *TREM2*, *CD9* – these genes are part of pathways involved in ECM reorganization. This expression signature has similarities to recently described studies in liver cirrhosis and pulmonary fibrosis where it was demonstrated to be a pro-fibrogenic phenotype (44, 59). *SPP1*-expressing macrophages have been demonstrated to play a role in promoting primary CRC. They have the potential to influence CD8 function by their role as an immune checkpoint ligand (60–62). This fibrogenic phenotype was accompanied by changes in genes controlling various metabolic pathways including glycolysis, lipid transport and sphingolipid synthesis resembling atherosclerotic foam cells (18). Macrophage metabolism influences their functional phenotype (63). Our findings provide metabolic targets that can be perturbed to further understand their biology in the context of the TME. The mCRC macrophages with alterations in lipid metabolism have been demonstrated to be associated with poor prognosis in cancer, including in mCRC (4, 24).

Fibroblasts and macrophages play a critical role in supporting the immunosuppressive TME, including the phenotype of T cell exclusion (64). We discovered fibroblasts specific to the TME with the potential to regulate ECM properties that can in turn promote tumor growth. Importantly, using an independent mCRC dataset we demonstrated that this fibroblast gene signature is linked to a worse clinical outcome and accompanied by reduced number of lymphocytes. This result is supported by recent studies in primary CRC, which identified positive correlation between fibroblasts and *SPP1*^+^ macrophages. Their presence was associated with poor survival and accompanied by reduced lymphocyte infiltration (65, 66).

The majority of mCRC tumors are MSS and unresponsive to T cell based immune checkpoint blockade. Hence, the gene expression programs and macrophage-fibroblast interactome represents potentially targetable elements in the TME of these patients. These targets are of interest also in other cancers to enable the modulation of the immunosuppressive stroma and improve immunotherapy response (64). In mouse models of cancer, TREM2 blockade resulted in TAM reprogramming and increased response to PD-1 immunotherapy (67). The CXCL12-CXCR4 interaction is also being investigated in clinical trials (52).

The gene expression programs we have discovered can potentially be influenced by tissue dissociation processes. We used the same dissociation protocol for matched normal liver to enable a controlled comparison between tumor and normal microenvironment lineages. This is reflected in the low number of hepatocytes we recovered (**Fig. 1B**), since adequate dissociation of normal liver requires specially developed dissociation protocols (17).

## Data Availability Statement

Sequencing data has been released under dbGAP identifier phs001818.v3.p1. Cellranger matrices will be available on https://dna-discovery.stanford.edu/research/datasets/.

## Code Availability Statement

Custom code used for CACTI and spatial proximity analysis is available at https://github.com/Kmason23/CACTI_Proximity_Test

## Authors’ contributions

AS, GP and HPJ designed the study. AS and BTL acquired the data. AS, BTL, KM, ZZ, NRZ developed the methodology. AS, KM, SMG, XB, ASu, XT, HJL, CJS, QN analyzed the data. AS, KM, QN, NRZ, HPJ interpreted the data. AS and HPJ wrote the manuscript with input from all authors. HPJ supervised the study.

## Acknowledgements

We are grateful to all patients who participated in the study as well as their families. We thank Christine Handy, Christina Wood-Bouwens and Alison Almeda for assistance in sample collection. Figures 1A, 5A, 6A were created using Biorender.com.

## SUPPLEMENTAL DATA

Supplemental Figures S1 – S6. Format: PDF

Supplemental Tables S1– S8. Format: XLSX

