## Supplemental figures for "Colorectal cancer metastases in the liver establish immunosuppressive spatial networking between tumor associated *SPP1*^+^ macrophages and fibroblasts"

#### LEGENDS TO SUPPLEMENTAL FIGURES

**Supplemental Figure 1.** (A) UMAP representation of dimensionally reduced data following batch-corrected graph-based clustering of all tumor epithelial cells annotated by sample. (B) Heatmap depicting expression of five highest significantly expressed genes (adjusted p-value < 0.05) per patient. (C) Heatmap representation of inferred single-cell CNV profiles of all tumors and reference cells.

**Supplemental Figure 2.** Heatmap depicting cell states from monocytes and macrophages identified using Ecotyper NMF analysis, annotated by condition and genes with highest fold change in each state.

**Supplemental Figure 3.** (A) UMAP representation of dimensionally reduced data following batch-corrected graph-based clustering of all lymphocyte lineage cells annotated by cell types. (B) Heatmap depicting average expression of selected genes for each cell type. (C) UMAP representation annotated by condition. (D) Heatmap depicting cell states from CD8 T cells identified using EcoTyper NMF analysis, annotated by condition. (E) Violin plot depicting the expression of cytotoxicity gene signature in normal and tumor CD8 T cells with T-test p-value.

**Supplemental Figure 4.** (A) Dot plot depicting expression levels of respective ligands and receptors together with the percentage of cells expressing them. (B) Top 10 percent of interactions inferred between cell types in the TME. Edge weights are proportional to the number of interactions, circle sizes are proportional to the number of cells in each group, edge color represents the cell type as sender. For scale, autocrine interactions in CAFs are 68.

**Supplemental Figure 5.** Graphical representation of cell types identified in CODEX analysis in image data from respective patients.

**Supplemental Figure 6.** (A-C) Comparisons with Pearson correlation between (A) proportions of cell lineages in samples with both scRNA-seq and CODEX data. (B) Average expression of LGALS3 and CD68 in macrophages across all patients. (C)  Average expression of COL4A1 and ACTA2 in CAFs across all patients.

#### Supplemental Figure 1

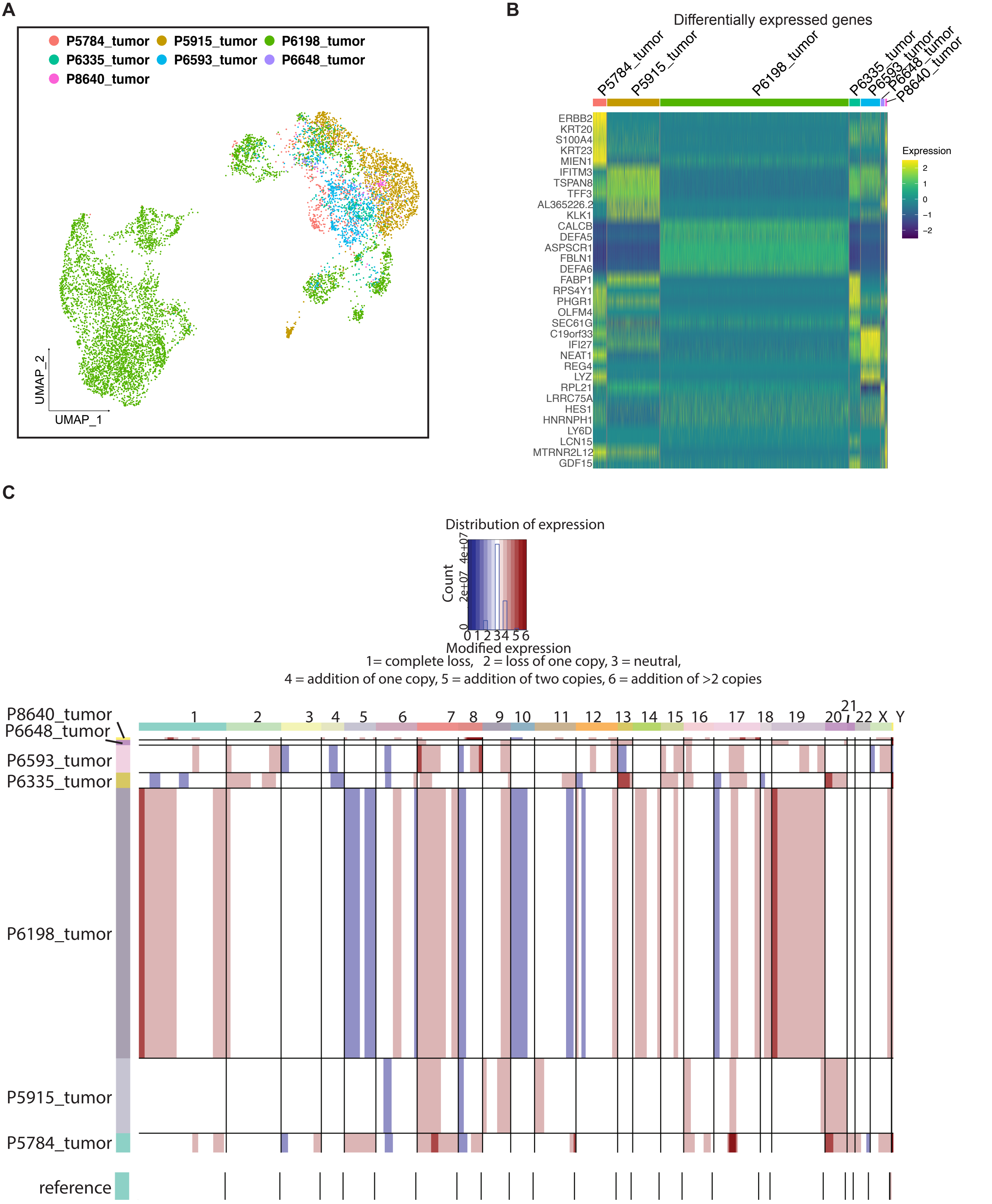

#### Supplemental Figure 2

**A**

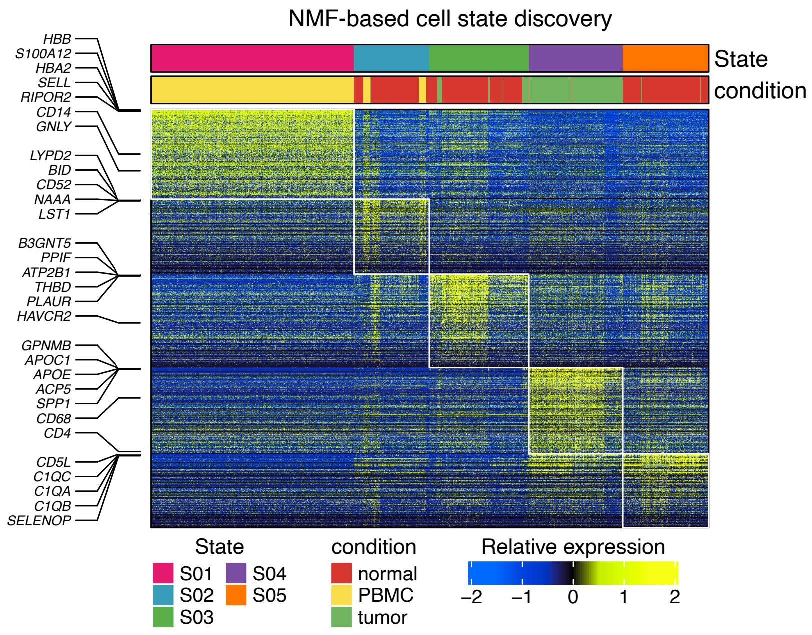

Supplemental Figure 3

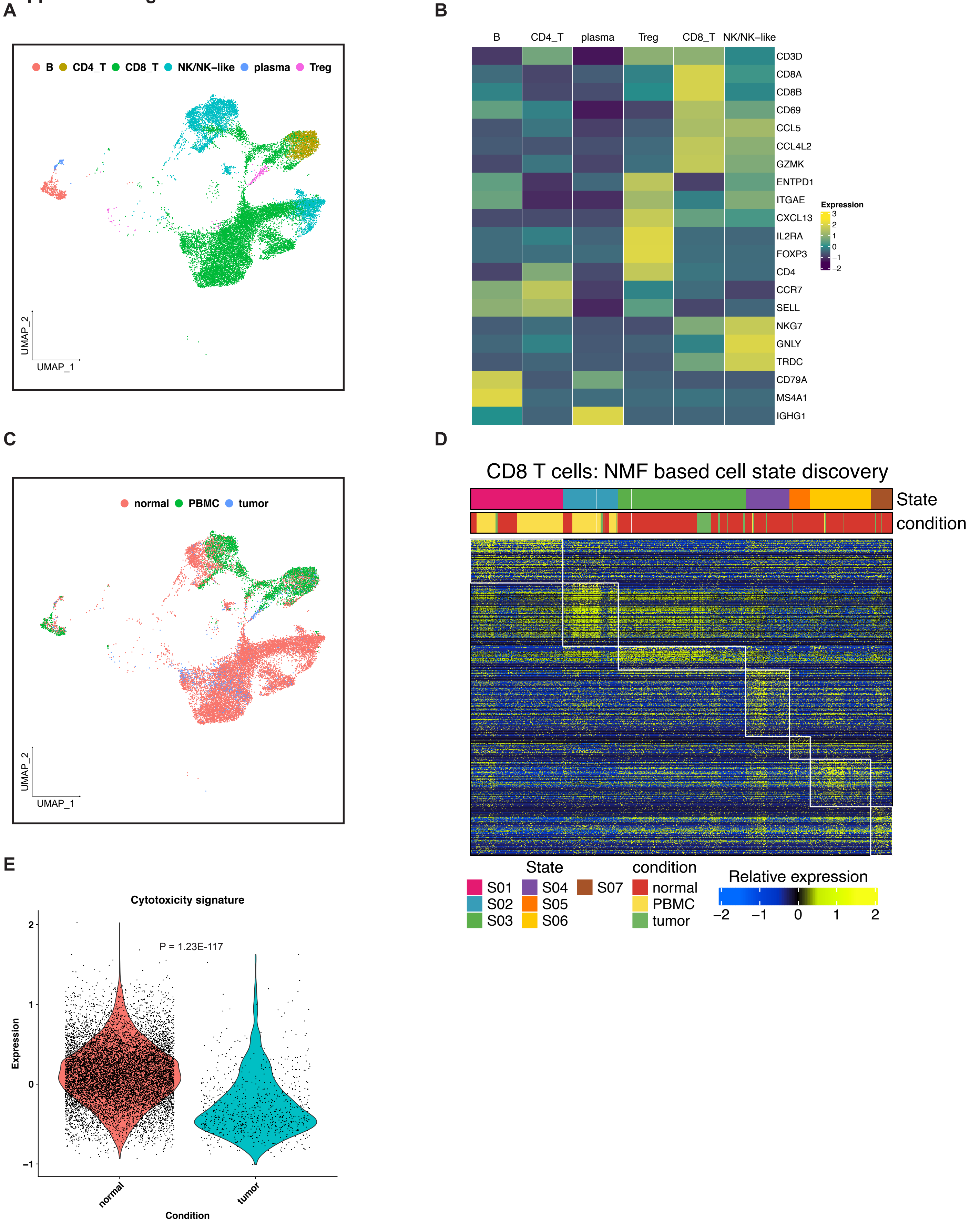

#### Supplemental Figure 4

A

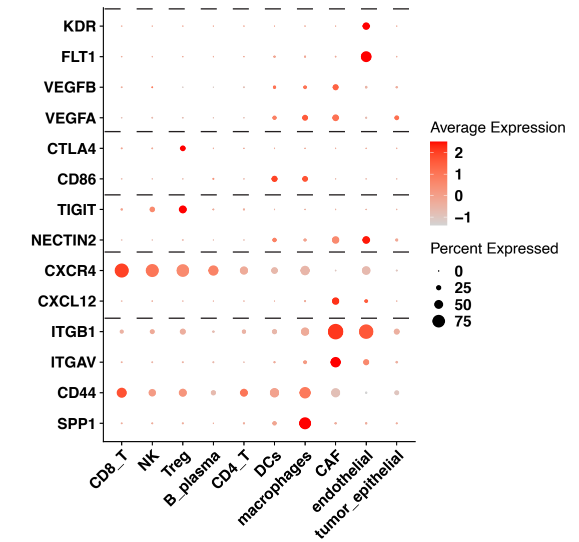


B

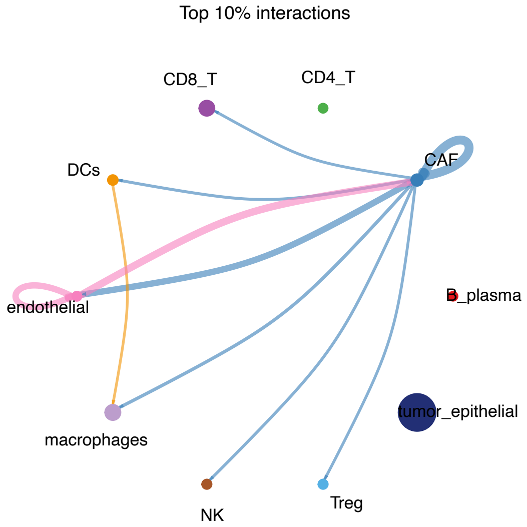

### Supplemental Figure 5

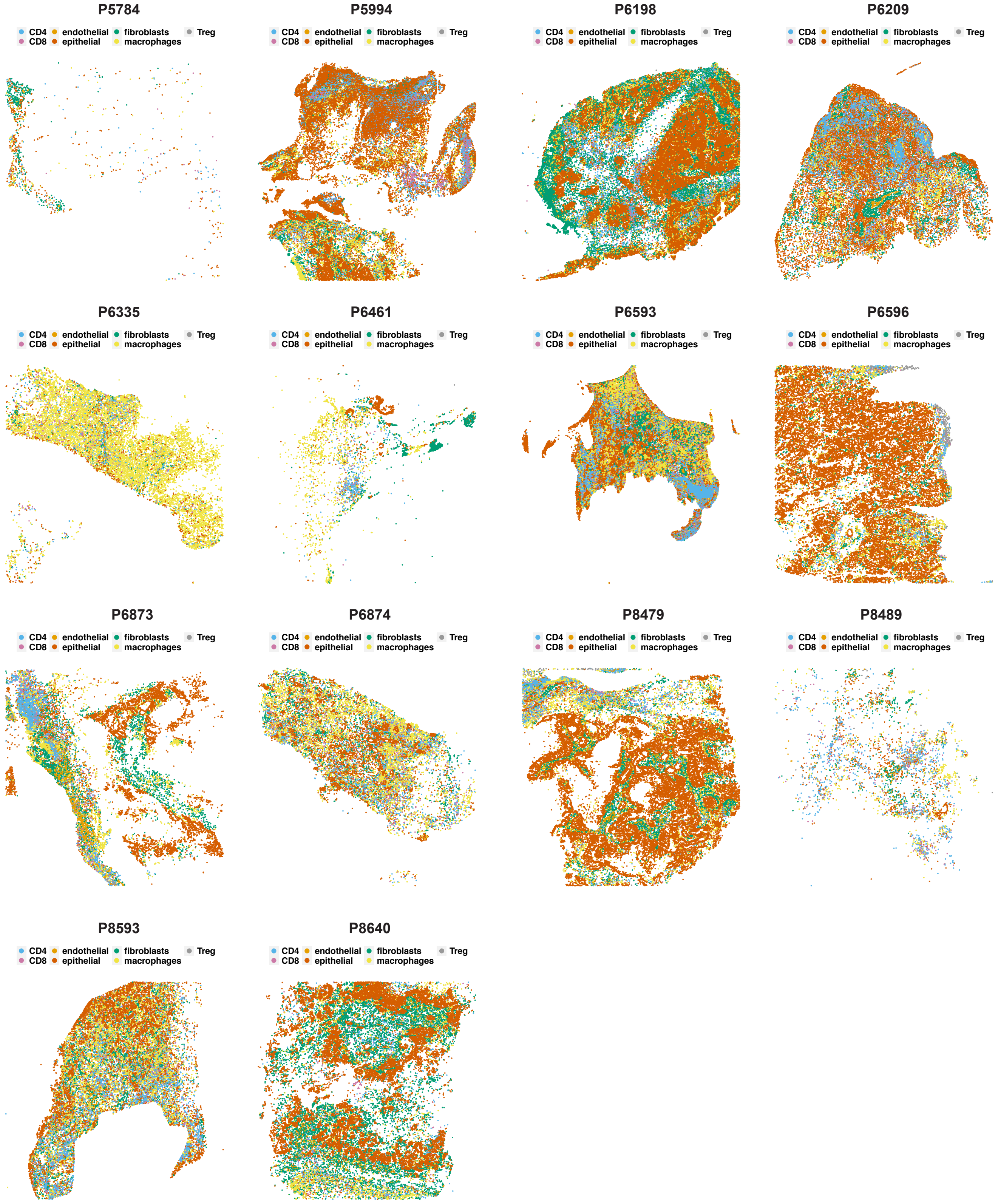

**Supplemental Figure 6**

**A**

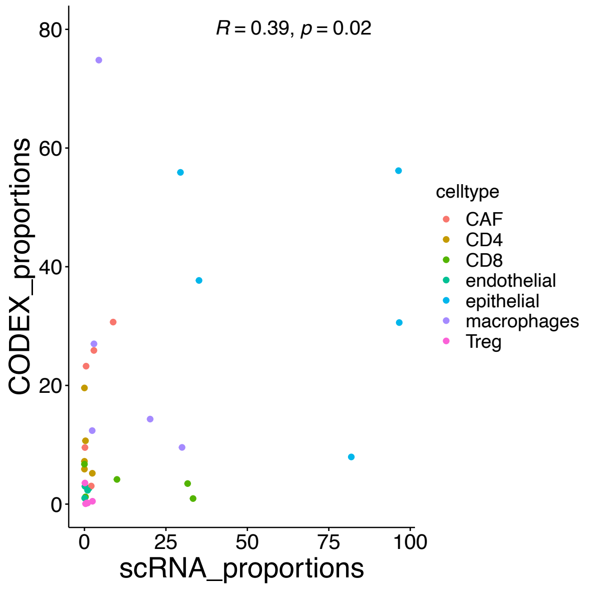


**B**

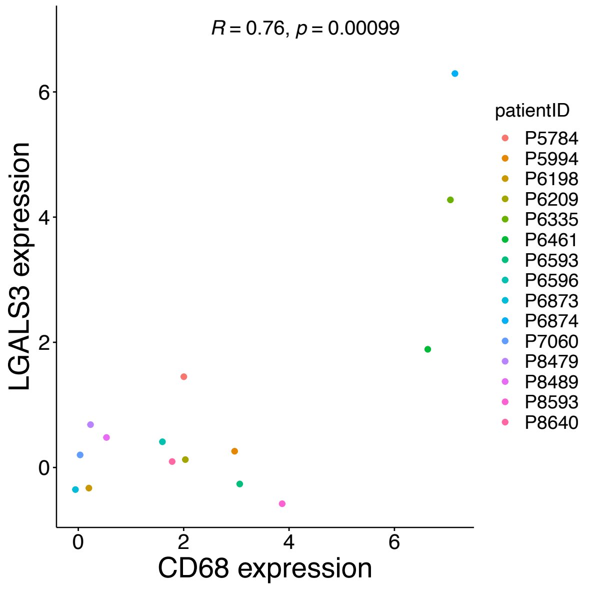


**C**

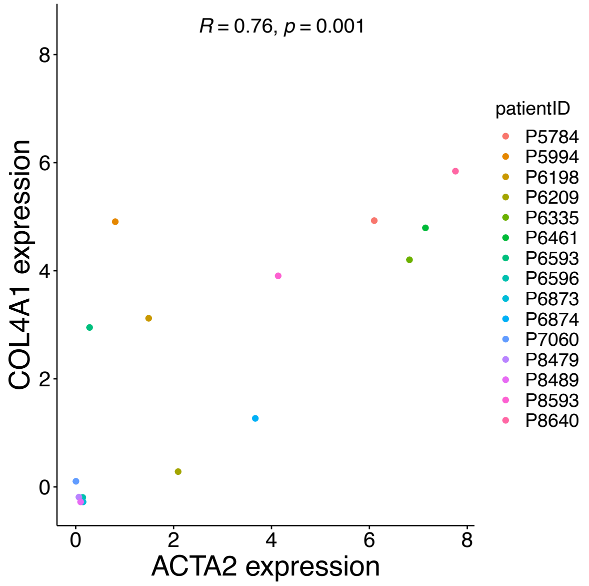
